# megaMine: a scalable, rule-based framework for mining gene-cancer-drug evidence from biomedical literature

**DOI:** 10.64898/2026.08.06.743392

**Authors:** Muhammad Junaid, Karolina Hanna Prazanowska, Ha-Eun Jeong, Yebin Ryu, Jeff Choi, Joon-Yong An, Su Bin Lim

**Author notes:** To whom correspondence should be addressed. Su Bin Lim, Department of Biochemistry & Molecular Biology, Ajou University School of Medicine, Suwon 16499, Korea.

## Abstract

The rapid expansion of the oncology literature has outpaced manual curation of clinically relevant gene-cancer-drug associations and oncogenic driver evidence. Existing automated approaches often lack transparency or are difficult to scale across heterogeneous data sources. To address this gap, we developed megaMine, a transparent, rule-based, and context-aware literature-mining framework that integrates therapeutic and driver evidence from PubMed, PubTator, and Europe PMC by combining entity recognition, hierarchical heuristics, and contextual labeling. In therapy mode, megaMine was applied to approximately 100,000 oncology articles published between 2015 and 2025, yielding more than 23,000 structured sentence-level evidence records, with standardized annotations for drug response, resistance, and study context. Internal evaluation of context labels showed the strong separability between efficacy and non-efficacy evidence using ridge logistic regression (AUROC = 0.915; AUPRC = 0.941). Benchmarking against NCI/OncoKB-supported drug-cancer associations showed that curated clinical associations had higher megaMine composite evidence scores than unlabeled comparison pairs [median (IQR): 25.6 (9.07-72.5) vs. 3.61 (1.69-8.69); Wilcoxon rank-sum test, P < 2.2 × 10^−16^]. In driver mode, megaMine retrieved mutation- and biomarker-related evidence from an *ERBB*-focused gastric cancer query, generating 750 evidence rows from 200 PMIDs. These results demonstrate that deterministic and interpretable approaches can support scalable evidence extraction for downstream applications such as knowledge graph construction and literature-based evidence synthesis.

## Introduction

The rapid expansion of genomic and oncology research has produced a vast amount of clinically relevant information embedded in unstructured biomedical text. Beyond patient records, millions of molecular biology and oncology publications describe complex relationships among genes, drugs, variants, cancer types, and therapeutic response. However, this wealth of textual information remains largely underutilized because automated extraction and normalization of contextual gene-cancer-drug evidence remains challenging . Traditional biomedical natural language processing has made substantial progress in clinical narrative mining, including the extraction of diagnosis, medication, procedures, and patient-level events from electronic health records . However, genomic oncology literature presents distinct extraction challenges due to complex combinations of gene symbols, sequence variants, cancer subtypes, therapeutic agents, response terms, and experimental or clinical context . These entities are highly ambiguous and require precise normalization, particularly for genomic variants that are frequently reported in non-standard formats across the literature . Therefore, deterministic rules and heuristic systems remain valuable for genomic literature mining because they offer interpretable, transparent, and reproducible evidence prioritization and sentence-level provenance, particularly when distinguishing true therapeutic or driver relationships from simple co-occurrence.

Several resources now provide programmatic access to large-scale biomedical literature and entity annotation. For example, the National Center for Biotechnology Information (NCBI) Entrez Programming Utilities (E-utilities), PubTator, and Europe PMC provide programmatic access to biomedical literature and entity annotations . However, these resources primarily support literary retrieval or entity-level annotation and do not directly convert heterogeneous annotations into structure and context-aware gene-cancer-drug relationships. Curated precision-oncology knowledge bases such as the NCI targeted drug therapy lists and OncoKB cover a fraction of the available evidence, focusing on selected biomarkers with regulations or guidelines while omitting broader metadata . Thus, an important gap remains: the lack of a scalable and reproducible framework that integrates the totality of biomedical evidence and converts it into structured, context-aware gene-cancer-drug evidence for translational decision support.

To address this gap, we developed megaMine, a scalable, rule-based framework that consolidates heterogeneous oncology literature into structured gene-cancer-drug evidence. The system operates in two complementary modes: 1) therapy mode extracts and normalizes therapeutic associations, including treatment response versus resistance, line of therapy, study design, and relevant clinical context; 2) driver mode identifies gene-cancer evidence related to oncogenic drivers, tumor suppressors, prognostic markers, and functional or mechanistic tumor assays. Together, these modes integrate clinical actionability and biological insight to support evidence-driven precision oncology. By prioritizing deterministic rules over opaque machine-learning models, megaMine ensures interpretability and reproducibility, enabling systematic integration of evidence for precision oncology.

## Methods

### Systematic overview

megaMine was implemented as a sequential modular Python workflow (>v3.9; Fig. 1) for extracting and structuring therapy- and driver-related evidence from oncology literature. The complete rule-library categories and representative patterns are summarized in Supplementary Table S1. The pipeline consisted of four stages:

1. Literature retrieval: PubMed abstracts and metadata were gathered via NCBI E-utilities, and full-text annotations were collected from Europe PMC.
2. Annotation integration: Entity information from PubTator and Europe PMC was harmonized and combined.
3. Heuristic extraction: Rule-based methods identified gene-drug-cancer connections and classified their biological context (e.g., efficacy, resistance, mutation, or toxicity).
4. Normalization and structuring: Extracted entities were linked to standard biomedical ontologies, producing analysis-ready, tabular outputs.

**Fig. 1.**
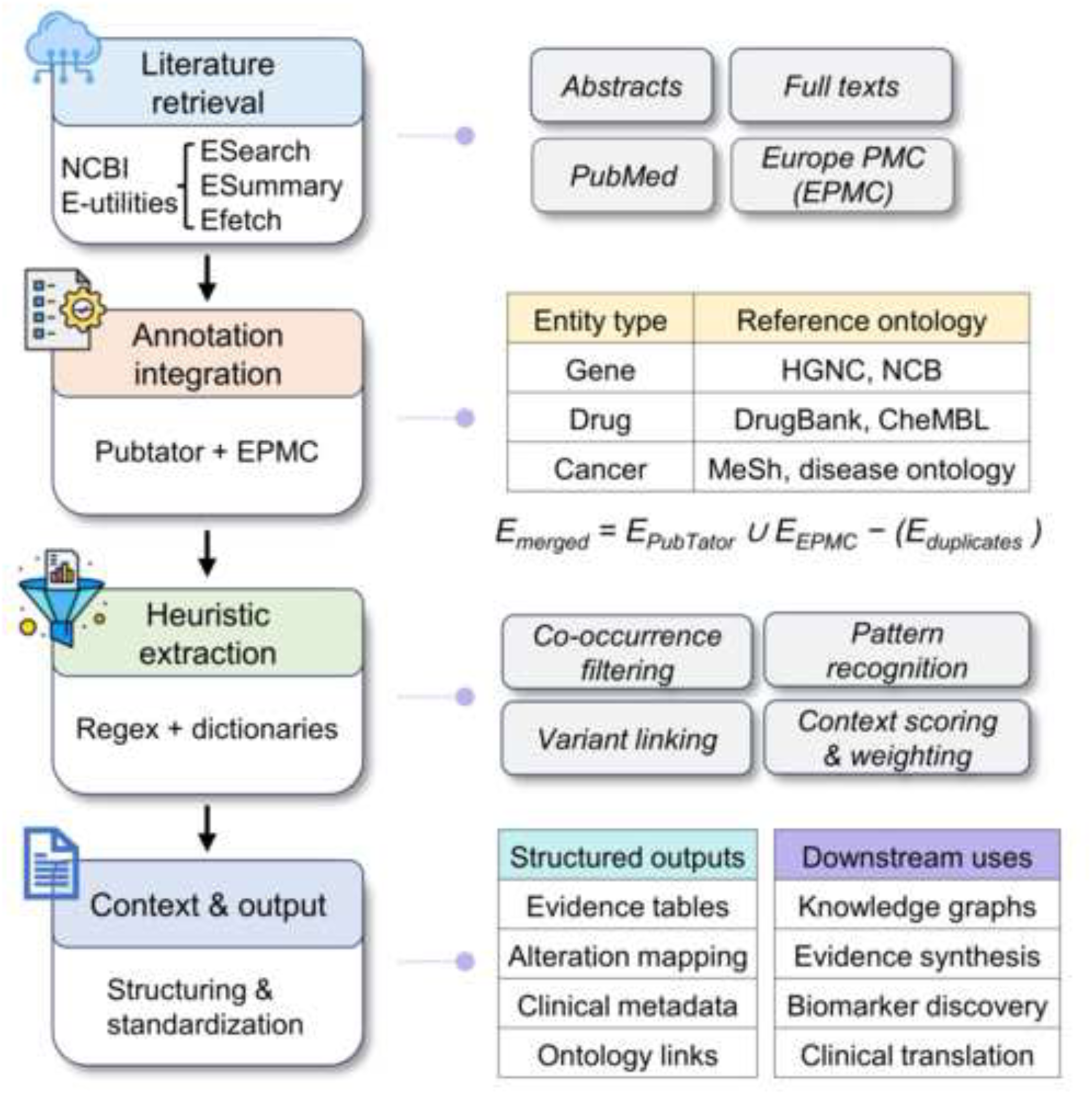
megaMine system overview. A four-stage pipeline mines pharmacogenomic evidence from literature.

### Literature retrieval

Articles were selected using oncology- and pharmacogenomics-focused PubMed queries combining cancer terms with gene, drug, mutation, therapy, inhibitors, antibody, resistance, and response-related keywords. Retrieval was restricted to publications from 2015 to 2025, and the therapy-mode analysis was capped at 100,000 PubMed records after PMID deduplication. PubMed abstracts were retrieved via NCBI E-utilities using *esearch.fcgi* (PMID discovery; with optional year-binned pagination to avoid clipping) and *esummary.fcgi* (metadata), and *efetch.fcgi* (XML abstracts) (titles, abstracts, and metadata). In addition, full-text annotations were obtained from Europe PMC through the annotations API if open-access articles are available. When a PMCID was available, and the --use-pmc-fulltext flag was enabled, sectioned full-text content was also retrieved via the Europe PMC REST service, focusing on open-access biomedical articles. Rate limiting was applied to ensure compliance with API policies (0.02 s delay per call; 0.12-0.34 s per batch, depending on NCBI key usage). The exact search strings, retrieval parameters, and capped-record selection are provided in Supplementary Table S2. This approach expanded the corpus beyond abstract-only content and enabled capturing contextual relationships across full-text sections.

### Annotation integration

megaMine constructed a unified, article-level annotation view by joining PubTator and Europe PMC records based on PMID and harmonizing their entity payloads prior to any rule-based extraction. PubTator documents were fetched in batches (≤150 PMIDs) and parsed to collect genes/proteins, diseases, chemicals (drugs), and mutations from passages.annotations, using infons.type to identify categories (e.g., gene, drug, disease, mutation). In parallel, EPMC annotations were retrieved via the Europe PMC REST service (SciLite) and parsed from the top-level annotation list, where entity types (e.g., Gene_Protein, Disease, Drug, Mutation) were mapped to a common internal schema. For each PMID, annotations from PubTator and Europe PMC were combined into a single unified set, and duplicates were removed based on normalized text forms (lowercase conversion, trimmed spaces, and punctuation removal) or shared identifiers. The source of each annotation (PubTator or EPMC; abstract or full text) was retained for subsequent weighting and quality control.

Furthermore, megaMine validated and normalized gene (biomarker) symbols using the HGNC public REST API, generating an in-memory set of approved symbols and discarding ambiguous or unapproved entries. Raw drug strings were detected with flexible text patterns (e.g., oncology-specific suffixes -nib, -mab, -parib, -ciclib, and “inhibitor(s)”) and then filtered against a curated oncology whitelist loaded via the --drug-whitelist option (ChEMBL, ChEBI, and DrugBank in plain-text or CSV format). Finally, cancer-related terms were standardized using canonical long-form/short-form mappings to ensure consistency in downstream heuristic processing.

### Heuristic extraction

After pre-annotation, the core therapy-based extraction module of megaMine identified gene-cancer-drug triplets within sentence boundaries using multi-stage heuristic rules. Each rule leverages lexical features (regular expressions and key phrases), positional cues (co-occurrence within a sentence), and semantic context labels derived from tokenized text. These rules were iteratively refined through empirical analysis of common linguistic patterns observed in oncology literature.

A sentence was considered eligible if it contained at least one HGNC-approved gene (G) validated through *GeneRegistry.is_valid(…)* and at least one whitelist-filtered drug (D) within the same sentence (s). This “same-sentence” constraint was strict, with no token-distance relaxation applied:

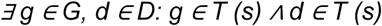

Sentences were assigned a single contextual label according to a hierarchical rule. Sentences expressing treatment outcomes (e.g., “treated with,” “responded to,” “sensitive to,” or quantitative metrics such as ORR, PFS, or OS) were labeled as efficacy. Those describing side effects (e.g., “adverse events,” “toxicity,” or specific cases like “hepatotoxicity”) were labeled as safety/toxicity. When the substring “toxicit” appears explicitly, the sentence was classified as toxicity; otherwise, it is labeled as safety. Sentences from articles indexed as reviews in PubMed were assigned the review label only if the sentence also contained review-related terms, such as “review”, “meta-analysis”, or “systematic”. All remaining sentences were labeled as background. Each contextual label is assigned to a priority weight:

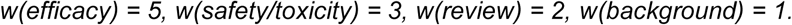

After labeling and weighting, megaMine retained only one key evidence sentence per primary section to prevent redundant counting. Section labels were standardized into four categories: Abstract, Methods, Results, and Discussion. For abstract-only PubMed records, structured abstract labels were mapped to these categories when available; otherwise, the entire abstract was treated as Abstract. Title and abstract text were both mapped to the Abstract category for evidence selection. Duplicate records within each PMID were collapsed when they shared the same gene, alteration, cancer type, section, evidence label, and sentence to ensure non-redundant results. For Europe PMC full-text records, available section headings, figure captions, and table captions were similarly mapped to the same primary section categories. Within each primary section, the sentence with the highest priority weight was retained. Efficacy evidence received the highest priority (5), followed by safety/toxicity (3) (Fig. 2A). The review evidence was assigned an intermediate low priority (2) because review or meta-analyses sentences may summarize prior evidence but are not treated as direct therapeutic observations. Background mentions received the lowest priority (1).

**Fig. 2.**
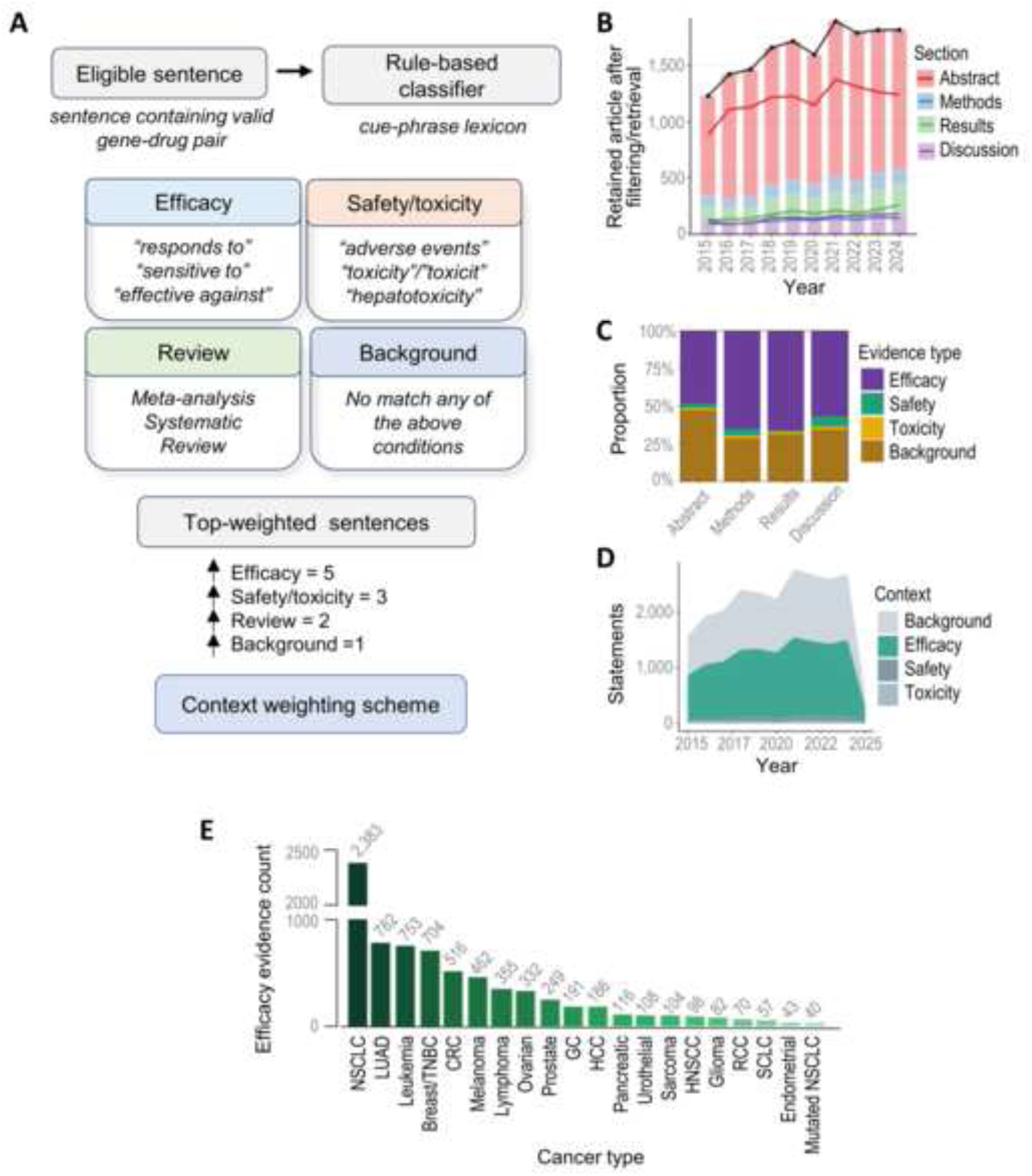
Context labeling via cue-phrase lexicons and evidence summary. (A) sentences are labeled by cue phrases and assigned a priority weight (w: efficacy = 5; safety/toxicity = 3; review = 2; background = 1). Within each section, the sentence with the highest weight is retained. (B) The number of papers retained per year after retrieval, annotation integration, and rule-based filtering. (C, D) Stacked bar plots show the mix of evidence types-efficacy, safety/toxicity, review, and background across primary sections, while the area plot illustrates how these evidence contexts have changed over time. (E) Top cancers by efficacy rows (normalized to canonical disease names). NSCLC: non-small cell lung cancer; SCLC: small cell lung cancer; LUAD: lung adenocarcinoma; CRC: colorectal cancer; TNBC: triple-negative breast cancer; EC: endometrial cancer; PCa: prostate cancer; PDAC: pancreatic ductal adenocarcinoma; PC: pancreatic cancer; RCC: renal cell carcinoma; UC: urothelial carcinoma; GC: gastric cancer; EAC: esophageal adenocarcinoma; ESCC: esophageal squamous cell carcinoma; HCC: hepatocellular carcinoma; CCA: cholangiocarcinoma; BTC: biliary tract cancer; GBM: glioblastoma; HNSCC: head & neck squamous cell carcinoma.

### Normalization and structuring

After selecting the highest-priority sentence in each section, megaMine attached an alteration only if it appears within the same sentence as the gene-drug pair (e.g., *G12D*, c.2573T>G, exon 19 deletion, or *EML4*-*ALK*). Fusion gene names were verified and standardized against HGNC-approved gene symbols and canonicalized (uppercase, suffix-stripped, partners sorted). If no alteration is detected, the record were retained as context-only. In megaMine, an evidence sentence refers to a retained sentence-level record as a structured output row, linking a gene, drug, and cancer type together with associated annotations, including alternation status, evidence context label, source section, PMID, and clinical metadata.

General study metadata were appended from the article-level text, including trial type or phase, treatment setting, stage or TNM classification, histology, omics details, PD-L1/TMB/MSI status, and reference genome version. The most frequently detected drug within each deduplicated evidence row was labeled as drug_primary, while co-administered agents are stored under combination_drugs. Deduplication was performed within each PMID without restricting the output to one evidence per article; multiple evidence rows could be retained from the same PMID when they differed by gene, alteration, cancer type, or evidence sentence. The therapy_type field was inferred from drug suffixes (e.g., -nib, -mab, -parib). Finally, the therapeutic_active flag is set to “yes” only for efficacy-labeled evidence rows.

### Oncogenic alteration extraction

In addition to the therapy-focused method, megaMine also includes a driver mode to identify oncogenic alterations independent of drug context. This module applies the biomarker-based heuristic rules to identify whether a gene alteration exhibits oncogenic, tumor-suppressive, or prognostic behavior in a specific cancer context. A sentence is considered eligible when it contains the normalized HGNC-validated gene symbol together with at least one alteration pattern or biomarker-context pattern keyword matched by megaMine rule library. The rule library includes patterns for amino-acid substitutions, nucleotide variants, exon events, gene fusions, copy-number alternation terminology, therapy-response context, resistance, study design, clinical setting, immune biomarkers, TMB/MSI status, and reference genome metadata. For driver-mode evaluation, we used a targeted gastric cancer query focused on *EGFR*/*ERBB*-family genes and oncogenic-activity terms, including driver, oncogene, amplification, overexpression, and copy-number alteration. The exact driver-mode query, filtering terms, and output are provided in Supplementary Table S3.

### Context-label consistency and benchmarking

To assess the internal consistency of megaMine’s rule-based context labels, we trained a ridge logistic regression classifier using TF-IDF features from the summary_sentence column. Efficacy-labeled records were treated as the positive class, and all other context labels were grouped as non-efficacy. The model was evaluated using 5-fold cross-validation, and performance was summarized using AUROC, AUPRC, and calibration analysis. This analysis assessed label consistency, not external extraction accuracy against a manually curated gold standard.

Second, to assess enrichment of clinically supported therapy relationships, literature-derived drug-cancer pairs were benchmarked against NCI targeted therapy listings and OncoKB therapeutic evidence levels 1– 3/R1. NCI/OncoKB-supported pairs were treated as reference-supported, while all others were used as an unlabeled comparison set. The megaMine composite evidence score and publication count were evaluated using ROC analysis, and AUROC values were compared to assess whether context-weighted scoring improved enrichment beyond raw publication frequency.

## Results

### Corpus scale and yield

We applied megaMine therapy mode to a capped set of 100,000 oncology-related PubMed records published between 2015 to 2025, yielding 23,808 structured evidence sentences. The final therapy-mode output is a compact tabular dataset, in which each row represents a retained evidence sentence linking a gene, drug, and cancer type, together with associated alteration when detected such as HGVS variant, exon, or gene fusion. Representative therapy-mode evidence records are shown in Supplementary Table S2, with the complete therapy-mode output retained in the full-output sheet of the same table.

### Evidence composition

The final retained dataset was enriched for efficacy-related evidence, reflecting the priority weighting used during sentence selection. Fig. 2B shows the annual number of retained articles after retrieval and filtering, with standardized evidence-section trends shown for Abstract, Methods, Results, and Discussion.

Across the standardized evidence sections, efficacy-related sentences constituted the majority of retained evidence (Fig. 2C). Safety and toxicity evidence represented a smaller subset of retained statements, consistent with the treatment-outcome focus of oncology literature. Over time, the number of retained efficacy-related statements generally increased together with the overall retained literature volume (Fig. 2D). The apparent decrease in 2025 likely reflects incomplete year coverage and the indexing at the time of retrieval rather than a biological or literature trend. Efficacy-related evidence sentences were most frequently extracted from studies on non–small cell lung cancer (NSCLC/LUAD), colorectal (CRC), breast/TNBC, melanoma, and leukemia studies. These cancer types dominate modern oncology literature and represent a major focus of drug development (Fig. 2E).

### Gene-drug landscape across diseases

The most commonly identified gene-drug pairs included several recognizable precision-oncology relationships, such as *EGFR*-osimertinib, *ALK*/*ROS1*-crizotinib, and *BRAF*-vemurafenib, which are also represented in curated oncology resources or supported by clinical literature. In context breakdowns, most gene-drug pairs originated from efficacy-labeled sentences highlighting patient’s response to treatment. In contrast, some also included safety or toxicity mentions, indicating potential adverse effects (Fig. 3A). Furthermore, therapy composition across different cancer types highlighted that targeted and immunotherapy approaches were most prevalent in NSCLC, LUAD, HCC (hepatocellular carcinoma), and leukemia. At the same time, chemotherapy predominated in prostate, RCC (renal cell carcinoma), and ovarian cancers (Fig. 3B).

**Fig. 3.**
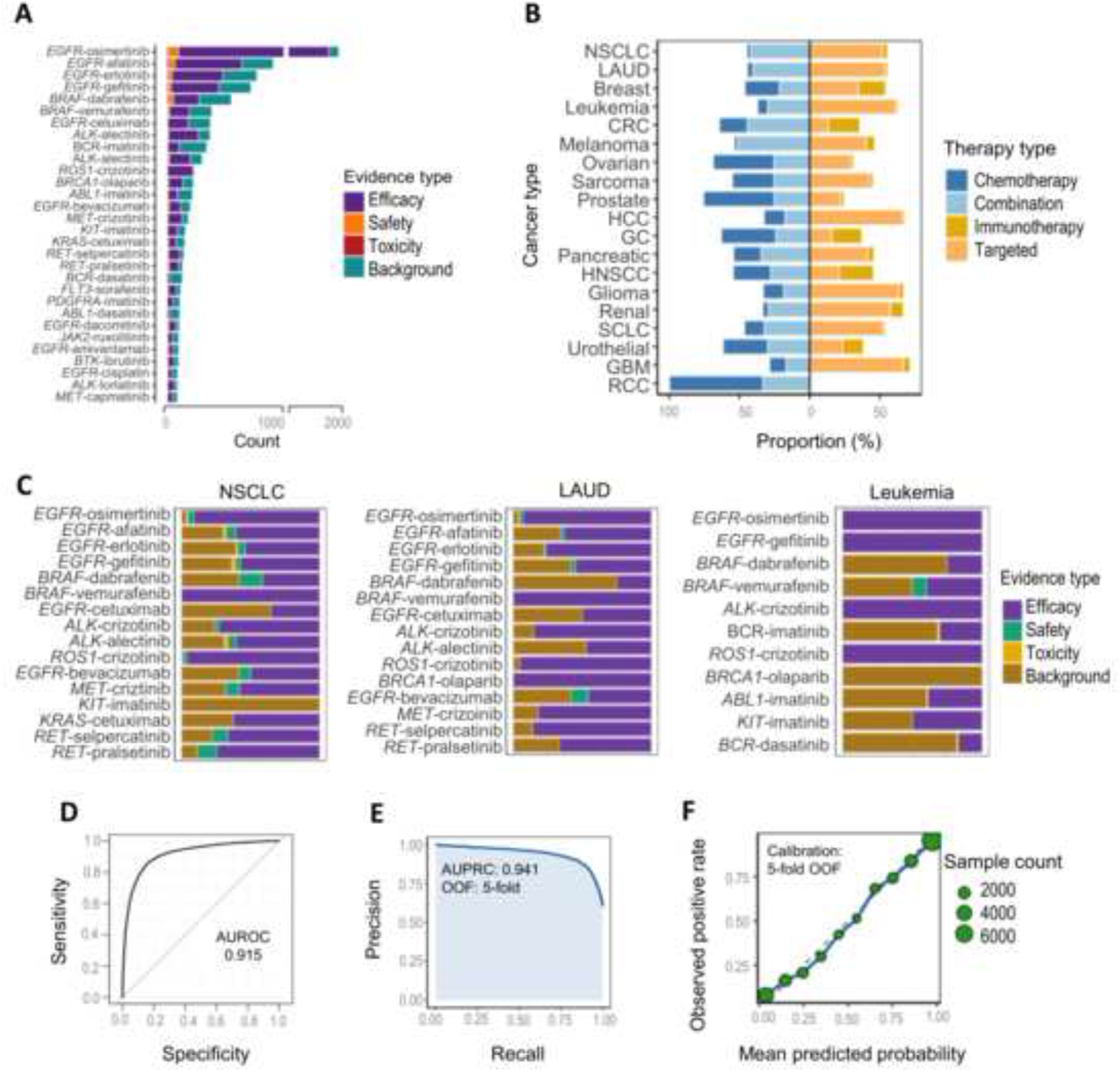
Therapy composition and model’s efficacy-label separability. (A) Top 30 gene-drug pairs showing how much of the evidence comes from efficacy, safety, toxicity, or background contexts. (B) Centered stacked bars show the composition of each therapy type across major cancers. (C) Gene-drug pairs with bars showing the proportion of evidence for each cancer, with stacked context compositions normalized within each disease to allow comparison of relative evidence patterns across cancers. (D) AUROC curve from 5-fold cross-validation (out-of-fold predictions) shows strong performance, with an AUC of 0.915 indicating excellent separation between efficacy and non-efficacy sentences. (E) Precision-recall curve from the 5 out-of-fold predictions shows strong performance, with a AUPRC of 0.941, indicating high precision across broad recall ranges despite class imbalance. (F) Calibration plot showing observed vs. predicted probabilities with curve closely follows the ideal line, indicating that the model predictions.

Contextual analysis across cancer types showed that efficacy-related evidence predominated across most disease studies, reflecting the literature’s focus on treatment outcomes (Fig. 3C). These results show that megaMine preserves both entity relationship and sentence-level evidence context, allowing gene-drug association to be stratified by cancer type and evidence class.

### Internal consistency of context labels

To assess the internal linguistic consistency of the megaMine’s rule-based context labels, we trained a ridge logistic regression model TF-IDF features derived from retained “summary_sentence” records.

The ROC curve showed strong separability between efficacy and non-efficacy sentences, with an AUROC of 0.915 (Fig. 3D). The precision-recall curve also showed strong performance, with an AUPRC of 0.941, indicating that efficacy-labeled sentences contained consistent lexical patterns even under class imbalance (Fig. 3E). In addition, to assess probability reliability, a calibration analysis was performed (Fig. 3F). The calibration curve showed that the predicted probabilities closely matched the observed positive rates, with most points lying near the ideal diagonal. Larger bubbles represent bins with more sentences and therefore provide more stable estimates. These findings further support the consistency of the megaMine’s deterministic context labeling.

### Therapy-mode benchmarking

Therapy-mode benchmarking showed that the megaMine composite evidence score distinguished NCI/OncoKB-supported drug-cancer from unlabeled comparison set with an AUCROC = 0.803 (Fig. 4A). We next compared two internal megaMine metrics, row-level composite evidence vs publication count, defining the number of PMIDs mentioning the same pair with the same NCI/OncoKB labels. The composite score showed a modestly higher AUROC than publication count alone (0.816 vs. 0.803), suggesting that context-weighted scoring may provide incremental information beyond raw literature frequency; however, the small difference should be interpreted cautiously (Fig. 4B). NCI/OncoKB-supported drug-cancer pairs had higher megaMine composite evidence scores than unlabeled comparison pairs [median (IQR): 25.6 (9.07-72.5) vs. 3.61 (1.69-8.69); Wilcoxon rank-sum test, P < 2.2 × 10⁻¹⁶] (Fig. 4C).

**Fig. 4.**
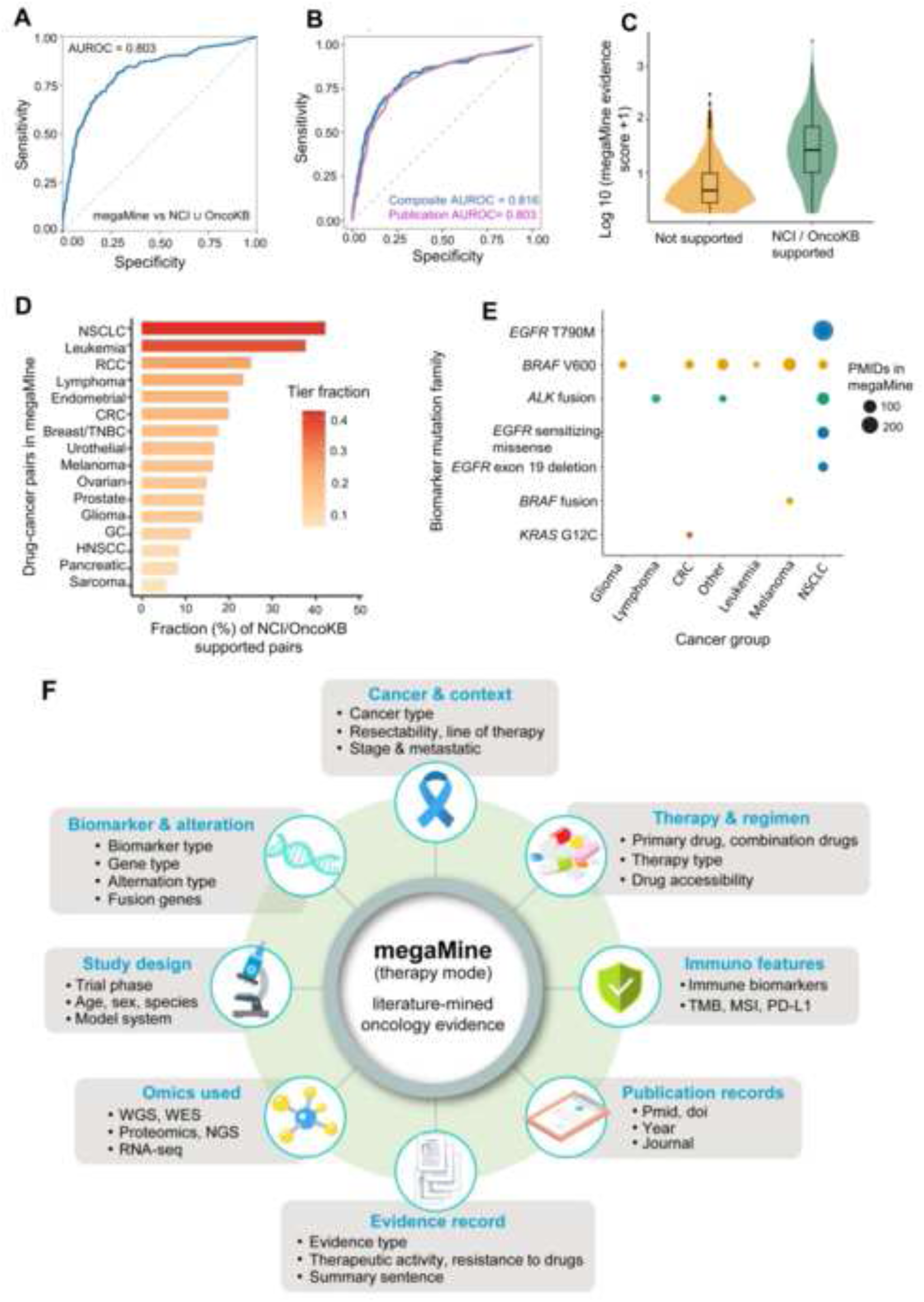
Validation of megaMine drug-gene-cancer and mutation associations against clinical knowledge bases. (A) ROC curve showing how well the megaMine literature evidence score distinguishes drug-cancer pairs that are supported by NCI approvals or OncoKB from all other pairs; the curve lies well above the diagonal random line, with an AUROC of 0.816, indicating strong enrichment of regulatory/guideline-supported therapies among high-scoring megaMine pairs. (B) ROC curves comparing two megaMine metrics: the full composite score (blue) vs simple publication count (pink), using the same NCI / OncoKB– supported vs. other drug-cancer labels as in panel A. (c) Violin/box plots showing log10-transformed megaMine evidence scores for drug–cancer pairs that are supported by NCI or OncoKB compared with all other pairs. (D) Bar plots showing the percentage of megaMine drug-cancer pairs on the x-axis that are supported by NCI or OncoKB. (E) Dot plot showing each point that represents a gene with a mutation family in a given cancer group, present in both megaMine and OncoKB. (F) megaMine therapy mode not only extracts biomarker-cancer-drug triplets (as in conventional clinical knowledgebases) but also a broader set of contextual features from each publication.

Cancer-type stratification showed that megaMine aligns most closely with curated knowledge for cancers where targeted therapy is well established. We observed the strongest overlap in NSCLC and leukemia, followed by renal cell carcinoma (RCC), lymphoma, endometrial cancer, colorectal cancer, and breast cancer (Fig. 4D). This pattern reflects the fact that these cancers have long-standing precision oncology programs with richer evidence bases. At the biomarker level, megaMine also recovered the key mutation families that define targeted therapy, such as *EGFR* exon 19 deletions, *EGFR T790M*, *BRAF* V600, *ALK* fusions, and *KRAS* G12C, each supported by many PMIDs in exactly the cancer types where they are clinically relevant (Fig. 4E). Finally, unlike curated resources such as NCI or OncoKB, which primarily catalogue validated drug–gene and cancer associations, the therapy mode of megaMine extracts a much broader range of contextual metadata from the literature (Fig. 4F).

### Driver-mode evidence extraction

After benchmarking the therapy mode, we evaluated driver-mode using a targeted query for gastric cancer and the *EGFR*/*ERBB* gene family. The research was capped at 1,000 papers and filtered using predefined oncogenic-activity terms, including driver, oncogene, amplification, overexpression, and copy number alteration. Details of the driver-mode query structure, filtering terms, and rule categories are provided in Supplementary Table S3. megaMine retrieved 750 evidence rows from 200 unique publications. The extracted records included recurrent driver-related contexts such as amplification, overexpression, copy-number alteration, oncogene terminology, and prognostic or mechanistic evidence. This analysis illustrates the broader utility of the same rule-based framework for organizing cancer-specific driver evidence, while formal benchmarking of driver-mode accuracy against manually curated driver annotations remains an important future step (Fig. 5).

**Fig. 5.**
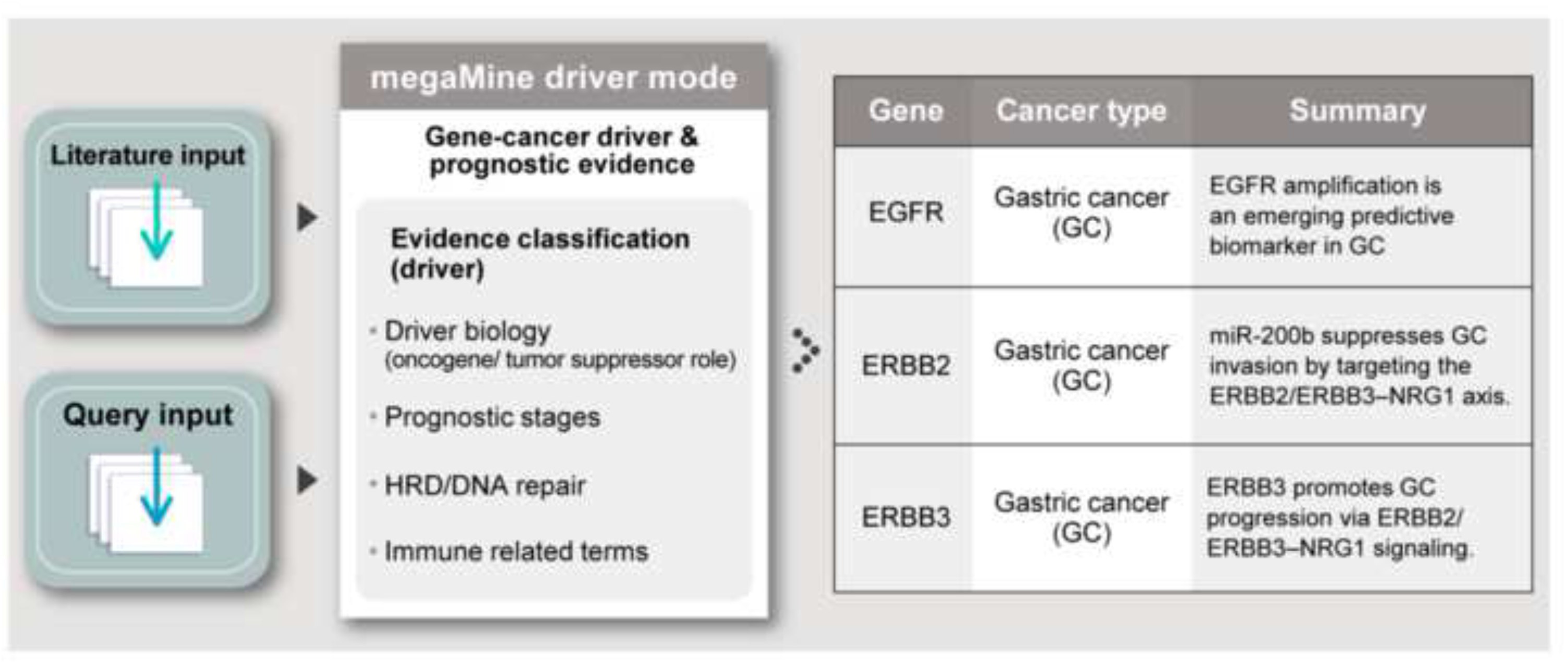
Driver-mode literature mining for gastric cancer WGS findings. megaMine driver mode classifier identifies gene-cancer drivers and prognostic evidence directly from biomedical literature using rule-based evidence tagging. The snapshot presents example evidence summaries supporting findings related to gastric cancer, with complete ranked evidence provided in Supplementary Table S3.

## Discussion

The rapid growth of oncology literature has created an urgent need for scalable systems that can convert heterogeneous biomedical text into structured, interpretable evidence. megaMine addresses this gap by integrating literature retrieval, entity annotation, rule-based contextual labeling, and evidence structuring into a unified framework for drug-gene-cancer and driver-related evidence mining. Our study demonstrates that a transparent, rule-based framework can extract clinically meaningful drug-gene-cancer relationships from large biomedical text collections while preserving sentence-level interpretability. The strongest therapy-mode signals corresponded to well-established precision oncology relationships. For example, megaMine recovered *EGFR*-Osimertinib evidence in NSCLC, consistent with the established role of Osimertinib in *EGFR-*mutant lung cancer . The framework also identified the *BRAF*-vemurafenib association in *BRAF-*mutant NSCLC and *ROS1*-crizotinib evidence in fusion-positive cancers, including lung cancer, ovarian cancer, and selected case-report context . These examples indicate that context-aware, sentence-level mining can prioritize recognizable therapeutic relationships while retaining the evidence provenance needed for downstream review.

Consistent with the objective, the overall retained evidence composition reflects megaMine’s prioritization of treatment-relevant sentences, with efficacy-labeled evidence forming the largest proportion and safety/toxicity, review, and background evidence contributing smaller subsets. Furthermore, the disease-specific panels showed biologically plausible cancer stratified patterns, with clinically recognizable therapy relationships appearing in relevant tumor contexts. In practice, the resulting evidence record can be directly used for curation triage (e.g., filtering efficacy statements in the Results section), safety monitoring, or stratified analyses by disease type, mutation class, or study characteristics such as trial phase, clinical setting, stage/TNM, and immune features.

Beyond therapy extraction, the driver mode extends the utility of megaMine by organizing oncogenic and biomarker-related evidence outside a direct drug-response context. For example, in the gastric cancer *EGFR*/*ERBB*-focused analysis, megaMine retrieved evidence related to amplification, overexpression, copy-number alteration, oncogenic activity, and prognostic relevance, illustrating how the framework can support cancer-specific driver evidence review. Together, therapy mode and driver mode provide a more complete picture of literature-supported biomarker roles.

There are some limitations in our approach. First, the rules are designed to be precise, even if that means not capturing every possible mention. As a result, statements expressed through negation, speculation, or across multiple sentences may sometimes be missed. Second limitation is variations in drug names (brand vs. generic) or newly approved therapies that can also escape the whitelist, and some rare cancer subtypes that may not be simplified by standard naming. Third, the coverage depends on the available literature, which mainly includes English-language papers and open-access full-text articles from Europe PMC. More broadly, megaMine is dependent on the quality and completeness of the underlying literature. Extracted associations may reflect preliminary, context-specific, or lower-evidence studies and should therefore be interpreted as literature-derived evidence rather than definitive biological and clinical conclusions. Finally, classifier analysis evaluated the internal consistency and linguistic separability of efficacy versus non-efficacy labels, rather than full extraction against a manually curated gold standard. Future benchmarking using manually annotated datasets will be important to evaluate entity recognition, relationship extraction, and context-label accuracy more directly.

Looking ahead, megaMine can be improved step by step. Future updates in heuristic extraction rules may include better handling of negation, speculation, and cross-sentence links, as well as smarter parsing of tabular content and figure captions. We also plan to add a ranking layer to prioritize evidence while keeping sentence-level transparency. Regular updates of HGNC genes, drug lists, and benchmark datasets will help maintain accuracy and expand coverage.

## Conclusion

In summary, megaMine demonstrates that a transparent, rule-based, and interpretable framework can extract relevant drug-gene-cancer relationships from large-scale biomedical literature at the sentence level without relying on opaque models. By integrating data from PubMed and Europe PMC resources, applying standardized gene and drug normalization, and focusing on context-labeled evidence sentences, the system generates compact, reproducible, and traceable evidence outputs. These results highlight the potential of deterministic approaches for scalable and interpretable evidence extraction in precision oncology.

## Data availability

The mined outputs generated in this study, together with the literature retrieval queries, run parameters, and rule-library details, are provided in supplementary tables S1-S3. Full-text articles are not redistributed; however, all extracted records retain PMID/PMCID provenance to support traceability. The source code and Python package are publicly available at GitHub (https://github.com/Junaid13913/megaMine) including documentation and reproducible scripts.

## Acknowledgements

This work was supported by the Korea Health Technology R&D Project through the Korea Health Industry Development Institute (KHIDI), funded by the Ministry of Health & Welfare, Republic of Korea (HR22C1734 and RS-2025-02310331), the National Research Foundation (NRF) of Korea (RS-2020-NR049588, RS-2020-NR046270, 2022R1C1C1004756), and the Korea-US Collaborative Cancer R&D Program funded by the Ministry of Health & Welfare, Republic of Korea (RS-2025-02223036).

## Author contribution

S.B.L., J.-Y.A., and M.J. conceptualized and designed the study. M.J. and S.B.L. developed the main code and drafted the manuscript. K.H.P., J.-Y.A., Y.R., and H.-E.J. contributed to data interpretation, validation, and code review. J.C. provided critical manuscript review and methodological feedback. All authors reviewed and approved the manuscript.

## Ethics declarations

### Ethics approval and consent to participate

Not applicable. This study did not involve human participants or clinical samples.

---

Supplementary Table S1

Supplementary Table S2

Supplementary Table S3

